# Context-dependent reconfiguration and spatial reorganisation of the miR-199a-5p/XIAP axis in high-grade serous ovarian cancer

**DOI:** 10.64898/2026.03.11.711019

**Authors:** Teresa Muñoz-Galdeano, David Reigada, Manuel Nieto-Díaz, Magda Palka Kotlowska, Lorena González Gea, María García Peña, Belén Santacruz, Guillermo Iglesias, Rodrigo M. Maza

## Abstract

**Background:** High-grade serous ovarian cancer (HGSOC) is a leading cause of gynaecological cancer mortality, largely because of late diagnosis and platinum resistance. Impaired apoptotic execution is a defining feature of resistant disease, but the regulatory interactions underlying this phenotype remain incompletely understood. The relationship between miR-199a-5p and the anti-apoptotic factor XIAP across cisplatin-sensitive and cisplatin-resistant states was the focus of this study.

**Methods:** We investigated the miR-199a-5p/XIAP axis in paired cisplatin-sensitive (A2780) and cisplatin-resistant (A2780cis) ovarian cancer models and in human ovarian tissue using multiplex fluorescence in situ hybridisation and immunofluorescence.

**Results:** miR-199a-5p did not show a uniform repressive relationship with XIAP across cellular states. In sensitive cells, miR-199a-5p was consistent with basal XIAP repression and enhanced apoptotic responses following cisplatin exposure. In resistant cells, this relationship was attenuated, suggesting uncoupling between XIAP and apoptotic responsiveness after cisplatin. In human FFPE specimens, spatial analysis identified tumour-associated reorganisation of this axis, with broader miR-199a-5p distribution, greater overlap with XIAP, and a distinct perinuclear pattern in HGSOC.

**Conclusions:** The miR-199a-5p/XIAP axis is context-dependent and altered in cisplatin-resistant ovarian cancer. These findings support spatially resolved analysis as a useful framework for identifying regulatory heterogeneity and tumour-associated phenotypes linked to platinum-resistant disease.

## INTRODUCTION

Ovarian cancer mortality is driven by relapse and platinum resistance, while robust predictive biomarkers remain limited. Platinum resistance is a multifactorial, context-dependent phenotype, particularly associated with poor outcome in primary platinum-resistant disease [1], and biomarker development is constrained by the difficulty of detecting clinically relevant biological changes early enough to guide treatment [2]. High-grade serous ovarian cancer (HGSOC) shows marked intratumoural heterogeneity, with spatially distinct subclones and microenvironmental niches shaping drug response and tumour evolution [3]. Spatial profiling has shown that clinically relevant response regions may differ transcriptionally despite similar histology, supporting spatial molecular analysis in HGSOC [4]. Apoptotic competence is central to platinum response. Resistance models implicate impaired apoptosis as a key mechanism of platinum failure, and inhibitor-of-apoptosis proteins (IAPs) suppress caspase activation while integrating survival signalling [5,6]. In ovarian cancer, XIAP has been linked to platinum-resistant phenotypes, supporting analysis of its regulation beyond abundance [5,7]. MicroRNAs add a further regulatory layer, but bulk measurements limit their interpretation. Although miRNA dysregulation is part of resistance-associated circuitry, bulk profiling cannot determine whether miRNA–target relationships are preserved or spatially/subcellularly reorganised [8]. Mature miRNAs can localise outside the cytoplasm, including the nucleus, where they contribute to gene regulation [9,10]. miR-199a-5p has been associated with tumour-suppressive functions in ovarian cancer and recurrence- or chemoresistance-related settings [11]. Additional studies support a role for miR-199a-5p across ovarian cancer microenvironmental contexts [12]. We previously identified XIAP as a direct miR-199a-5p target in a mammalian experimental system [13], supporting examination of this interaction in ovarian cancer.

Here, we investigated whether the miR-199a-5p/XIAP axis contributes to cisplatin response and whether this relationship changes in resistant states. We combined CLIP-informed in silico prediction, functional analysis in paired cisplatin-sensitive and cisplatin-resistant ovarian cancer models and spatially resolved miRNA–protein mapping in human FFPE tissue. This strategy evaluates the miR-199a-5p/XIAP relationship as a context-dependent regulatory interaction whose spatial organisation may be relevant to platinum resistance in HGSOC.

## MATERIALS AND METHODS

Extended methodological details, including manufacturer information, culture conditions, antibody specifications and DIG-labelled probe sequences, are provided in the Supplementary Information.

### In silico miRNA–mRNA interaction analysis

Accessibility of the miR-199a-5p binding site within the human XIAP 3′ untranslated region (3′UTR) was assessed using STarMir [14], with the mature hsa-miR-199a-5p sequence, the human XIAP 3′UTR, the human V-CLIP-based model and 3′UTR-restricted settings. Outputs included hybridisation energy (ΔG_hybrid), seed-pairing configuration, accessibility features and the composite LogitProb value. Local RNA secondary structure around the predicted binding site was analysed using RNAfold. Human and murine XIAP 3′UTRs were examined under identical conditions to assess structural conservation.

### Patient samples

FFPE ovarian tissue samples were obtained from the Department of Obstetrics and Gynaecology, Hospital Universitario de Torrejón (Madrid, Spain). The cohort comprised HGSOC samples (N = 10) and non-neoplastic ovarian controls (N = 10) from surgery for benign gynaecological conditions. Representative haematoxylin and eosin-stained sections were reviewed by a certified pathologist to confirm diagnosis. For each FFPE block, a 3 mm punch biopsy was taken from the selected region of interest. Samples were anonymised, coded and analysed blinded.

### Cell culture

Human ovarian carcinoma A2780 cells and their cisplatin-resistant counterpart A2780cis were obtained from the Cell Bank of the Centro de Instrumentación Científica, University of Granada (Granada, Spain). Cells were STR-authenticated and confirmed free of bacterial, fungal and mycoplasma contamination. Culture media and supplements for each cell line are listed in Supplementary Table S2.

### Transfection and cisplatin treatment

Transient transfections used DharmaFECT-4 according to the manufacturer’s instructions. Cells were transfected for 24 h with 50 nM miRIDIAN hsa-miR-199a-5p mimic or non-targeting negative control mimic and, where indicated, treated with 12.5 μM cisplatin for a further 24 h. Control cells were maintained in cisplatin-free medium.

### RNA extraction and RT–qPCR

Total RNA was isolated from untreated A2780 and A2780cis cells using QIAzol Lysis Reagent and the miRNeasy Mini Kit. RNA concentration and purity were assessed using a NanoDrop ND-1000 spectrophotometer. For miR-199a-5p quantification, 10 ng of total RNA was reverse transcribed and amplified using a TaqMan MicroRNA assay, with U6 as endogenous control. For XIAP mRNA, 1 μg of total RNA was DNase I-treated, reverse transcribed and analysed using TaqMan chemistry, with 18S as endogenous control. Amplification was performed on a TaqMan 7900HT Fast Real-Time PCR System. Relative expression was calculated using the 2^-ΔΔCt method [15], with error propagation as described by Headrick [16]. Experiments included three independent biological replicates and three technical replicates per condition.

### Protein extraction and Western blot analysis

Proteins were extracted from A2780 and A2780cis cells under basal conditions or after transfection and cisplatin treatment. Cells were lysed in RIPA buffer with protease inhibitors, and protein concentration was determined by BCA assay. Equal protein amounts were separated by SDS–PAGE and transferred onto PVDF membranes. Membranes were blocked, incubated overnight with primary antibodies and then with HRP-conjugated secondary antibodies. Proteins were visualised by chemiluminescence under non-saturating conditions, and band intensities were quantified using ImageJ. Antibodies are listed in Supplementary Table S3. Biological replicate numbers are indicated in the corresponding figure legends.

### Fluorescence in situ hybridisation/immunofluorescence

miR-199a-5p detection in FFPE ovarian tissue sections was performed using an adaptation of the protocol described by Søe et al. [17]. Tissue sections were deparaffinised, rehydrated, subjected to antigen retrieval and proteolytic digestion, and hybridised with DIG-labelled LNA probes specific for miR-199a-5p or the negative control cel-miR-67. Probe sequences are listed in Supplementary Table S4. FISH signal was detected using alkaline phosphatase-conjugated anti-digoxigenin Fab fragments, followed by XIAP immunofluorescence and DAPI counterstaining. Images were acquired using an Olympus IX83 microscope. For cultured cells, A2780 and A2780cis cells grown on glass coverslips were processed under RNase-free conditions using an adapted DIG-labelled LNA-FISH protocol. Cells were permeabilised, acetylated, hybridised with miR-199a-5p or negative control probes, and sequentially processed for anti-digoxigenin detection, XIAP immunofluorescence and DAPI counterstaining. Images were acquired using a Leica DM5000B epifluorescence microscope. No signal was detected in negative control conditions.

### Cell viability and cell death assays

Cell viability was assessed by MTT assay in A2780 and A2780cis cells after transfection and cisplatin treatment. Late cell death was evaluated by propidium iodide (PI) staining as a measure of plasma membrane integrity. Effector caspase activation was assessed using CellEvent™ Caspase-3/7 Green Detection Reagent. In image-based assays, PI-positive or caspase-3/7-positive cells were calculated relative to total nuclei.

### Image and colocalisation analysis

Quantitative image analysis of basal XIAP and miR-199a-5p fluorescence, PI staining and caspase-3/7 activity was performed using QuPath v0.4.3. In cultured cells, nuclei were detected from DAPI fluorescence and relevant channel intensities measured. Basal fluorescence measurements included 556 A2780 and 564 A2780cis cells across three independent experiments. For PI and caspase-3/7 assays, approximately 1,500 cells per condition were analysed across five non-overlapping fields per replicate in four independent experiments. For tissue FISH/IF, ten regions of interest (ROIs; 750 × 750 μm) were analysed per sample, and XIAP and miR-199a-5p fluorescence intensities were quantified per detected object. ROI-level data were exported for downstream analysis. Colocalisation metrics were computed blinded in R (≥4.2) within the tidyverse framework. Channel-specific thresholds were defined using the 25th percentile of per-cell intensity distributions. Pearson correlation coefficients, Manders’ overlap coefficients (M1 and M2), and QuPath-derived cross-correlation metrics (CCF_MAX, CCF_MIN, K1, K2 and FWHM) were calculated from per-cell intensity data. ROI-level data were aggregated to one mean value per image for downstream statistical analysis.

### Statistical analysis

Statistical analysis was performed using GraphPad Prism and R (≥4.2). Data are expressed as mean ± standard error of the mean (SEM) unless otherwise stated. Linear mixed-effects models were used for functional assays, including MTT, PI, caspase-3/7 activation and XIAP protein levels under treatment, with transfection status, cisplatin treatment and their interaction as fixed effects and experimental day as a random intercept. For basal two-group comparisons, Student’s t-test, Welch’s t-test or Mann–Whitney U test were used as appropriate. A paired one-tailed t-test was used for basal XIAP immunoblot comparisons after miR-199a-5p transfection, based on a pre-specified directional hypothesis. Principal component analysis explored multivariate structure in colocalisation data. Tests are detailed in the figure legends and/or Supplementary Information. Statistical significance was defined as P < 0.05.

## RESULTS

### XIAP is a conserved target of miR-199a-5p

In our previous study [13], miR-199a-5p was identified among seven microRNAs predicted to target the rat XIAP 3′UTR and functionally validated as a direct regulator in a mammalian system. To assess whether this interaction was conserved in human ovarian cancer, we examined target-site accessibility and duplex stability. PITA analysis [27] showed canonical seed pairing and favourable hybridisation energies for miR-199a-5p within the human and mouse XIAP 3′UTRs (ΔG_hybrid = −15.4 and −15.2 kcal/mol, respectively; Supplementary Fig. S1 A,B). STarMir [14], a CLIP-informed prediction framework incorporating experimentally mapped miRNA–mRNA interactions [18,19], identified an accessible miR-199a-5p binding site in the human XIAP 3′UTR, with a LogitProb value above the functional threshold (>0.5; 0.543) (Supplementary Fig. S1 C). RNAfold modelling supported structural accessibility of the predicted binding region (Supplementary Fig. S1 D). Together, these data support a thermodynamically favourable and evolutionarily conserved miR-199a-5p/XIAP interaction, providing a rationale to examine its functional behaviour in human ovarian cancer models.

### Basal characterisation of the miR-199a-5p/XIAP axis in cisplatin-sensitive and cisplatin-resistant ovarian cancer cells

To determine whether basal alterations in the miR-199a-5p/XIAP axis were associated with cisplatin resistance, we compared transcript and protein expression in A2780 and A2780cis cells. TaqMan-based RT–qPCR showed significantly lower miR-199a-5p expression in A2780cis cells than in A2780 cells (fold change −2.58; t∟ = 3.46, p = 0.013; n = 3; Fig. 1A, upper panel), whereas XIAP mRNA did not differ significantly between cell lines (t∟ = 1.106, p = 0.165; n = 3; Fig. 1A, lower panel). Consistently, immunoblotting of whole-cell lysates from three independent experiments showed no significant difference in basal XIAP protein abundance between A2780cis and A2780 cells (1.52 ± 0.41 vs. 1.27 ± 0.31; t∟ = 1.709, p = 0.11; one-tailed paired t-test; 95% CI, −0.861 to 0.3; Fig. 1B).

**Figure 1.**
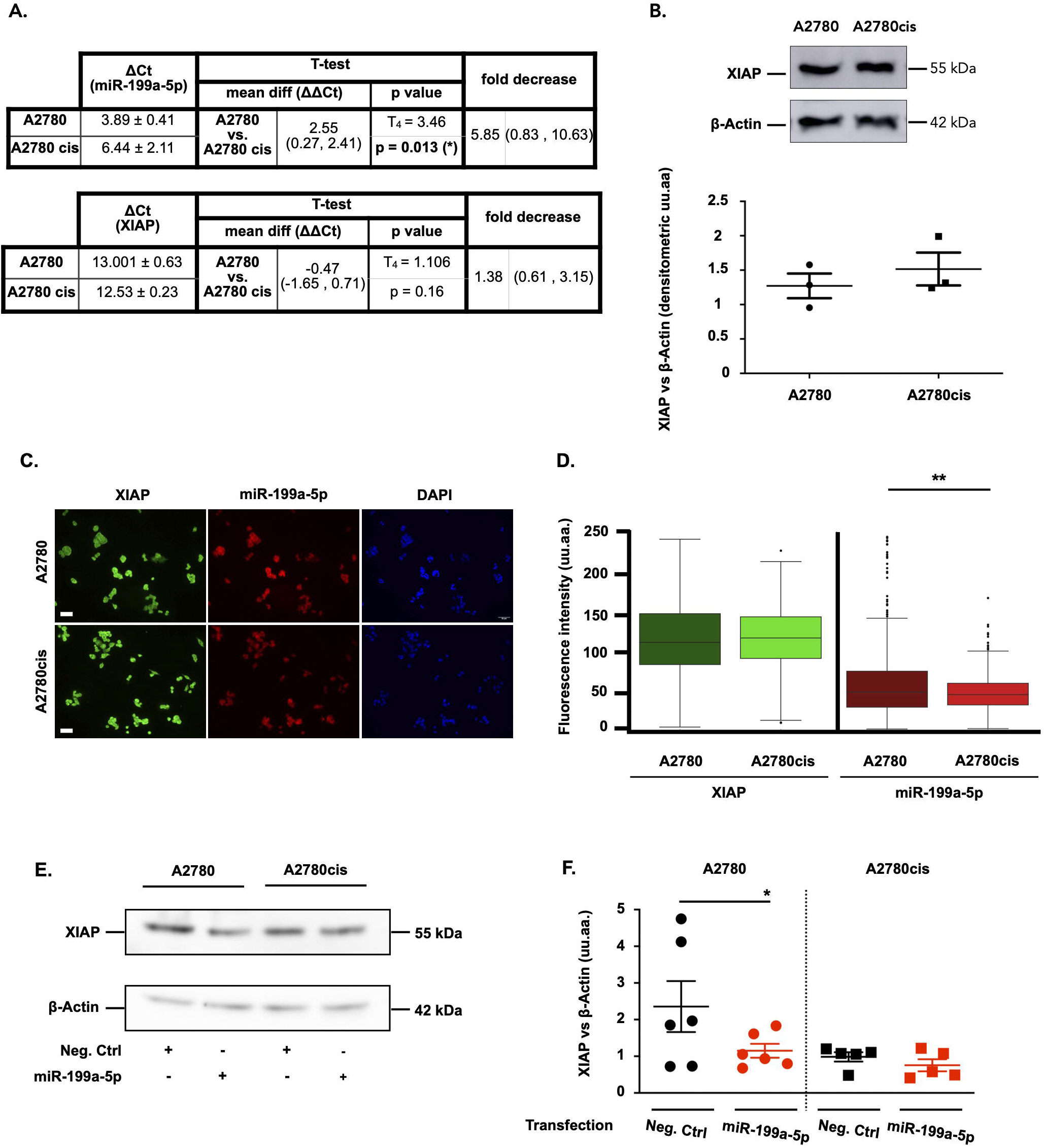
Basal expression of miR-199a-5p and XIAP in cisplatin-sensitive (A2780) and cisplatin-resistant (A2780cis) ovarian cancer cells. (A) RT–qPCR analysis of basal miR-199a-5p (upper panel) and XIAP mRNA (lower panel) expression in A2780 and A2780cis cells. Data are shown as ΔCt values, mean differences (ΔΔCt), p values and fold change estimates from Student’s t-tests. (B) Representative immunoblot showing basal XIAP and β-actin expression in A2780 and A2780cis cells (left), and densitometric quantification of XIAP normalised to β-actin (right). Data represent mean ± SEM from three independent experiments. (C) Combined fluorescence in situ hybridisation (FISH) for miR-199a-5p (red) and immunofluorescence for XIAP (green) in A2780 (upper panel) and A2780cis (lower panel) cells, with nuclei counterstained with DAPI (blue). Boxplots show per-cell fluorescence intensity distributions for each marker. Statistical analysis was performed using Mann–Whitney U tests. Scale bar, 50 μm. (E) Representative Western blot of XIAP in whole-cell lysates from A2780 (left) and A2780cis (right) cells transfected with either a negative control mimic or miR-199a-5p mimic under basal conditions. β-Actin was used as a loading control. (F) Densitometric quantification of XIAP normalised to β-actin from six independent experiments in A2780 cells (circles) and five independent experiments in A2780cis cells (squares). Statistical analysis was performed using one-tailed paired *t*-tests. *p* < 0.05.

We next assessed whether this pattern was reproduced at single-cell level. Combined fluorescence in situ hybridisation for miR-199a-5p and immunofluorescence for XIAP, performed across three independent experiments, showed reduced miR-199a-5p-associated fluorescence in A2780cis cells relative to A2780 cells (41.2 ± 21.7 vs. 60.5 ± 27.9 a.u.; p = 0.009; Fig. 1C,D), whereas XIAP-associated fluorescence did not differ significantly between cell lines (102.8 ± 44.0 vs. 91.2 ± 27.3; p = 0.209; Mann–Whitney U test). Thus, bulk and cell-associated measurements consistently identified reduced basal miR-199a-5p in the resistant phenotype without a significant basal increase in XIAP.

Finally, to determine whether miR-199a-5p retained basal repressive activity over XIAP, XIAP protein levels were analysed after miR-199a-5p transfection under untreated conditions (Fig. 1E). In A2780 cells, miR-199a-5p significantly reduced XIAP relative to the negative control (one-tailed paired t-test, p = 0.0359, n = 6; Fig. 1F, left). By contrast, no significant reduction was detected in A2780cis cells (p = 0.0765, n = 5; Fig. 1F, right). These findings indicate that cisplatin resistance is associated not only with reduced basal miR-199a-5p expression, but also with attenuated miR-199a-5p-dependent basal repression of XIAP.

### Effect of miR-199a-5p on cisplatin response in ovarian cancer cells

To examine whether miR-199a-5p modulates cisplatin response, we assessed cell viability, membrane integrity, apoptotic activity and XIAP protein levels in A2780 and A2780cis cells. First, the resistant phenotype of A2780cis cells was confirmed by MTT assays. A2780cis cells showed a higher cisplatin EC∟∟ than parental A2780 cells (44.56 μM vs. 12.16 μM; Supplementary Fig. S2), consistent with reduced cisplatin sensitivity.

Cell viability was then assessed after miR-199a-5p or negative control transfection, followed by treatment with or without cisplatin (Fig. 2). In A2780 cells (Fig. 2, left), the model identified significant main effects of cisplatin (F(1,7) = 15.29, p = 0.0058) and miR-199a-5p (F(1,7) = 10.66, p = 0.0138), with no significant interaction between both factors (F(1,6) = 0.714, p = 0.4305), indicating additive rather than interaction-dependent effects on metabolic viability. Consistent with the cisplatin main effect, cisplatin reduced viability across transfection conditions, reaching significance both in negative control-transfected cells (0.4213 ± 0.0343 vs. 0.4891 ± 0.0307; p = 0.0234) and in miR-199a-5p-transfected cells (0.3294 ± 0.0389 vs. 0.4357 ± 0.0213; p = 0.0234). miR-199a-5p alone showed a non-significant trend towards reduced viability relative to untreated controls (p = 0.0531). By contrast, in A2780cis cells (Fig. 2, right), neither miR-199a-5p transfection (F(1,7) = 0.435, p = 0.5305), cisplatin treatment (F(1,7) = 3.397, p = 0.1079), nor their interaction (F(1,6) = 0.0045, p = 0.9485) significantly altered viability. Together, these data support a context-dependent effect of miR-199a-5p on cisplatin-associated viability, evident in A2780 cells but not retained in A2780cis cells.

**Figure 2.**
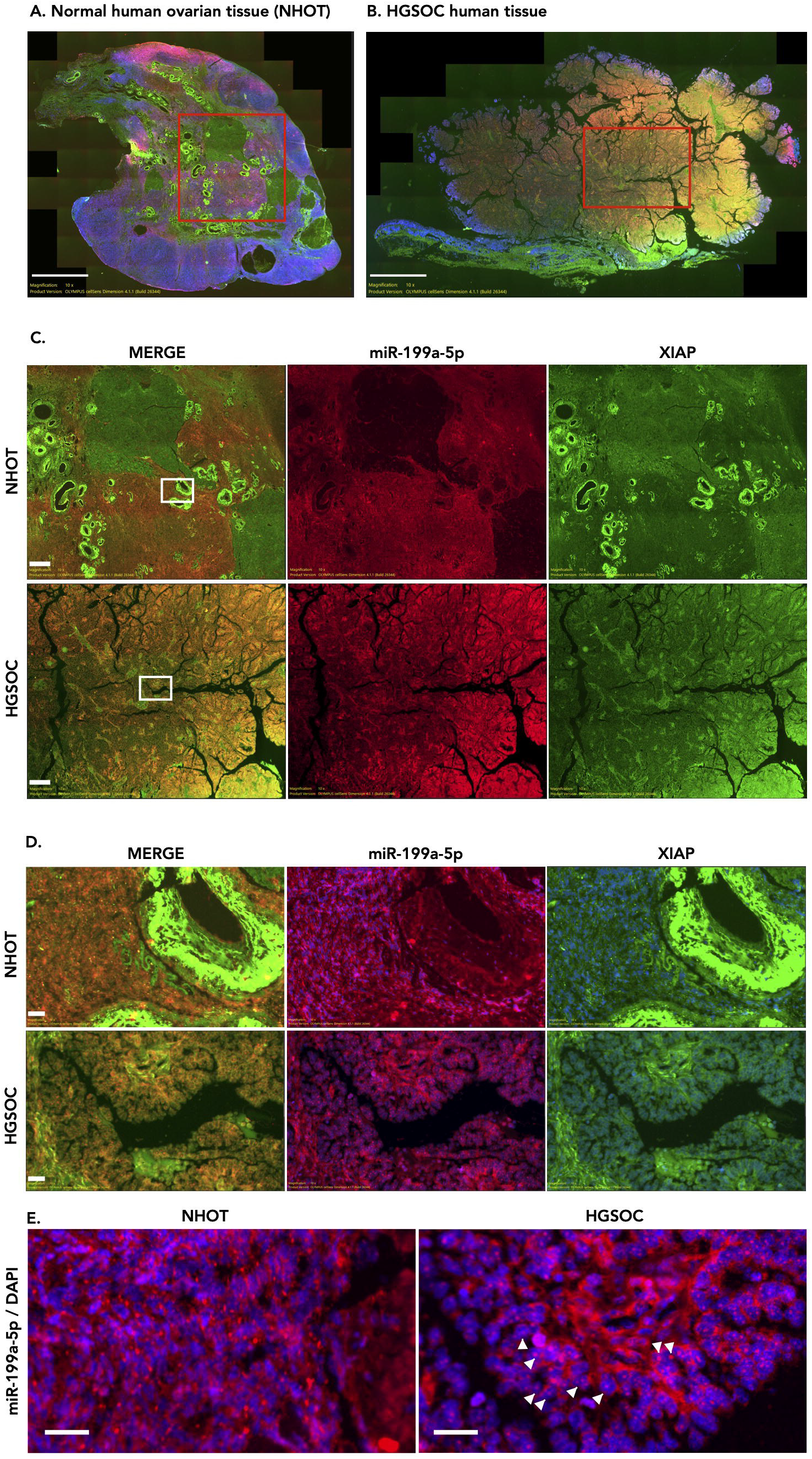
Effect of miR-199a-5p and cisplatin on cell viability in A2780 and A2780cis cells. A2780 (circles) and A2780cis (squares) ovarian cancer cells were transfected with either a negative control mimic (NC) or a miR-199a-5p mimic, and subsequently treated with or without 12.5 μM cisplatin (cisPt) for 24 h. Cell viability was assessed by MTT assay and expressed as optical density (OD∟∟∟–OD∟∟∟). Each point represents one independent experiment (n = 3), and bars indicate mean ± SEM. Filled symbols indicate untreated conditions (cisPt−), and open symbols indicate cisplatin-treated conditions (cisPt+). Statistical analysis was performed using a linear mixed-effects model followed by Tukey-adjusted post hoc comparisons. *p < 0.05.

### miR-199a-5p potentiates cisplatin-associated late cell death selectively in cisplatin-sensitive cells

Membrane integrity was next examined by propidium iodide staining to distinguish reduced metabolic viability from late cell death (Fig. 3). In A2780 cells (Fig. 3C, left), PI incorporation was influenced by miR-199a-5p transfection (F(1,9) = 7.6558, p = 0.0219) and by the miR-199a-5p × cisplatin interaction (F(1,9) = 6.7081, p = 0.0292), whereas cisplatin alone did not show a significant main effect (F(1,9) = 2.5721, p = 0.1432). Post hoc testing showed that the combined treatment increased PI-positive cells relative to cisplatin alone (t(9) = −3.788, p = 0.0185), while comparisons with untreated controls (p = 0.0524) and miR-199a-5p alone (p = 0.0633) remained just above the significance threshold. This pattern indicates that miR-199a-5p strengthened cisplatin-associated late cell death in sensitive cells rather than acting as an effective inducer alone.

**Figure 3.**
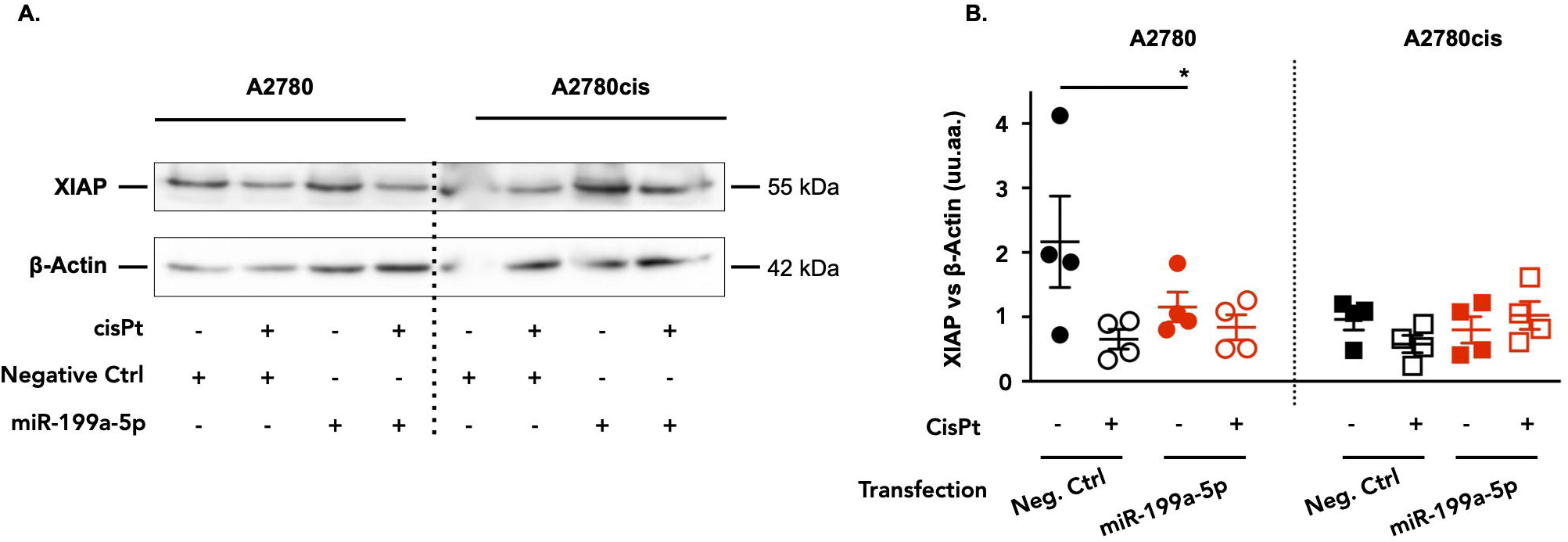
miR-199a-5p enhances cisplatin-associated cell death in ovarian cancer cells. (A, B) Representative phase-contrast and epifluorescence images showing propidium iodide (PI)-positive cells (red) in A2780 (A) and A2780cis (B) cells transfected with either a negative control mimic or a miR-199a-5p mimic and treated with or without 12.5 μM cisplatin (CisPt) for 24 h. (C) Quantification of PI-positive cells (%). Each point represents one independent experiment (n = 4), and bars indicate mean ± SEM. Statistical analysis was performed using a linear mixed-effects model with transfection, cisplatin treatment and their interaction as fixed effects, and experimental day as a random intercept, followed by Tukey-adjusted post hoc comparisons. *p < 0.05.

In A2780cis cells (Fig. 3C, right), PI incorporation remained stable across conditions, with no significant effect of miR-199a-5p (F(1,9) = 0.1223, p = 0.7346), cisplatin (F(1,9) = 4.0582, p = 0.0748) or their interaction (F(1,9) = 3.0564, p = 0.1144). The absence of a clear PI response in the resistant line suggested that any pro-death effect of miR-199a-5p might instead emerge at the level of apoptotic signalling rather than terminal membrane breakdown.

### Caspase-3/7 activation reveals a treatment-dependent pro-apoptotic effect of miR-199a-5p in A2780cis cells

To determine whether miR-199a-5p affected apoptotic signalling upstream of terminal membrane permeabilisation, caspase-3/7 activity was quantified (Fig. 4). In A2780 cells (Fig. 4B, left), caspase activation was driven mainly by cisplatin exposure (F(1,12) = 27.8896, p = 0.00019), whereas neither miR-199a-5p transfection (F(1,12) = 3.2912, p = 0.0947) nor the miR-199a-5p × cisplatin interaction (F(1,12) = 0.6172, p = 0.4473) reached significance. Post hoc analysis confirmed that cisplatin increased caspase-positive cells in both control-transfected cells (t(9) = −3.179, p = 0.0459) and miR-199a-5p-transfected cells (t(9) = −4.290, p = 0.0090), without significant differences between transfection conditions. Thus, the apoptotic response of sensitive cells was largely attributable to cisplatin.

**Figure 4.**
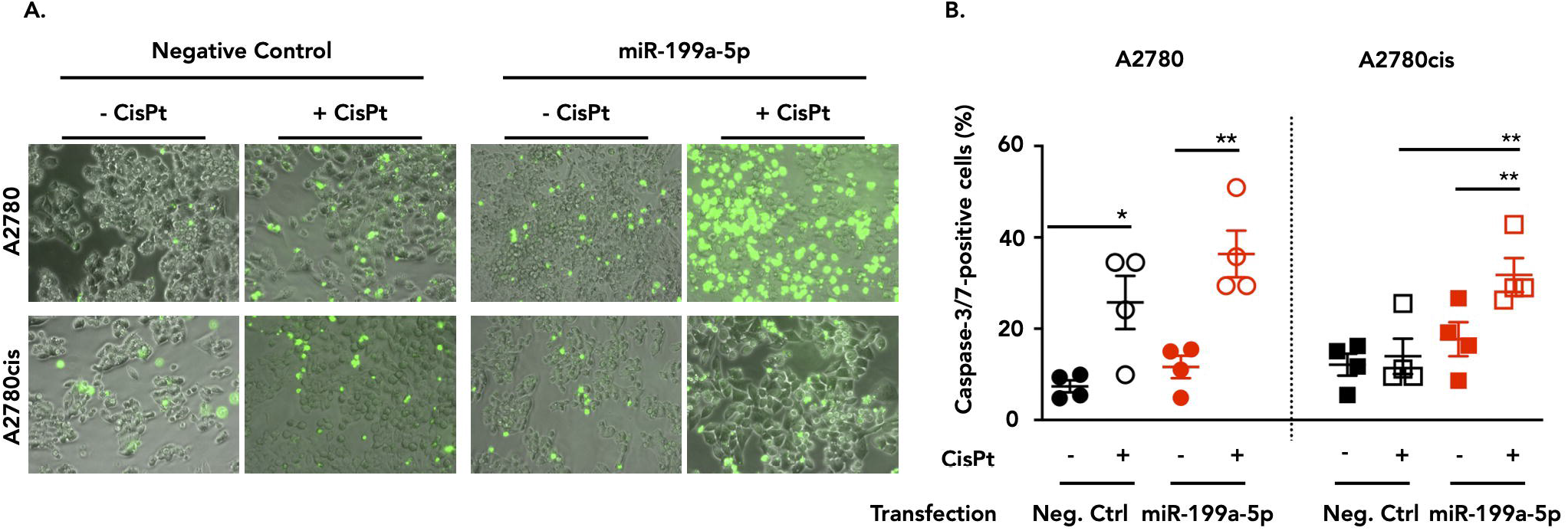
miR-199a-5p enhances cisplatin-associated caspase-3/7 activation in ovarian cancer cells. (A) Representative fluorescence microscopy images showing caspase-3/7-positive cells (green) in A2780 and A2780cis ovarian cancer cells transfected with either a negative control mimic or a miR-199a-5p mimic, and treated with or without 12.5 μM cisplatin (CisPt) for 24 h. (B) Quantification of caspase-3/7-positive cells (%). A2780 cells are shown on the left (circles) and A2780cis cells on the right (squares). Each point represents one independent experiment (n = 4), and bars indicate mean ± SD. Statistical analysis was performed using a linear mixed-effects model with transfection, cisplatin treatment and their interaction as fixed effects, and experimental day as a random intercept, followed by Tukey-adjusted post hoc comparisons. *p < 0.05, **p < 0.01.

**Figure 5.**
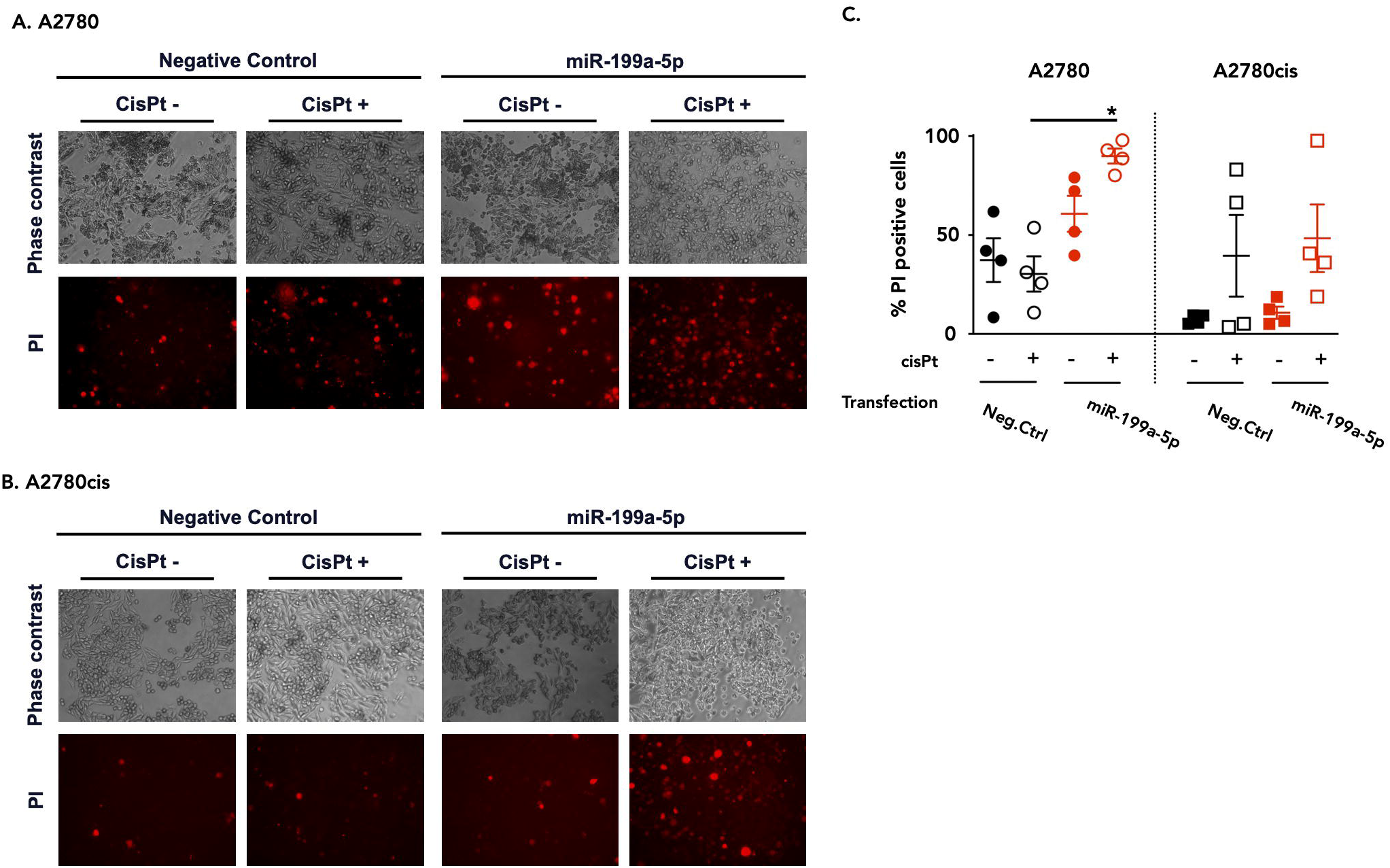
Effects of cisplatin and miR-199a-5p on XIAP protein levels in ovarian cancer cells. (A) Representative western blot analysis of XIAP in whole-cell lysates from A2780 (left) and A2780cis (right) ovarian cancer cells transfected with either a negative control mimic or a miR-199a-5p mimic and treated with or without cisplatin (CisPt; 12.5 μM, 24 h). β-Actin was used as a loading control. (B) Densitometric quantification of XIAP normalised to β-actin. A2780 cells are shown on the left (circles) and A2780cis cells on the right (squares). Each point represents one independent experiment (n = 4), and bars indicate mean ± SEM. Statistical analysis was performed using a linear mixed-effects model with transfection, cisplatin treatment and their interaction as fixed effects, and experimental day as a random intercept, followed by Tukey-adjusted post hoc comparisons. *p < 0.05.

A different pattern emerged in A2780cis cells (Fig. 4B, right), where cisplatin (F(1,9) = 11.4521, p = 0.0081), miR-199a-5p transfection (F(1,9) = 25.5065, p = 0.00069) and the miR-199a-5p × cisplatin interaction (F(1,9) = 6.9905, p = 0.0267) all contributed significantly to caspase-3/7 activation. Under cisplatin exposure, miR-199a-5p markedly increased caspase activity relative to the corresponding negative control (t(9) = −5.441, p = 0.0019), and cisplatin increased caspase activity only in miR-199a-5p-transfected cells (t(9) = −4.262, p = 0.0093). No transfection-dependent difference was observed without cisplatin, and cisplatin alone did not differ from untreated controls in negative control-transfected cells. These findings indicate that miR-199a-5p retains the capacity to modulate cisplatin-associated apoptotic signalling in A2780cis cells, even when this effect is not translated into reduced metabolic viability.

### Cisplatin exposure alters the miR-199a-5p/XIAP relationship in resistant cells

To determine whether caspase-3/7 activation was accompanied by downstream executioner caspase processing, cleaved caspase-3 was analysed by immunoblotting in A2780 cells (Supplementary Fig. S3 A) and A2780cis cells (Supplementary Fig. S3 B). In A2780 cells (Supplementary Fig. 3C, left), cleaved caspase-3 accumulation was significantly influenced by cisplatin (F(1,9) = 25.28, p = 0.0007), with no detectable effect of miR-199a-5p transfection (F(1,9) = 0.008, p = 0.931) or the miR-199a-5p × cisplatin interaction (F(1,9) = 2.76, p = 0.131). Post hoc analysis showed a marked cisplatin-induced increase in control-transfected cells relative to untreated controls (estimated difference = 1.201 ± 0.239, p = 0.0032). A similar directional increase was observed after miR-199a-5p transfection, although it did not remain significant after multiple-testing correction (estimated difference = 0.640 ± 0.239, p = 0.0973). In A2780cis cells (Supplementary Fig. S3 C, right), none of the model terms reached statistical significance (cisplatin: F(1,8) = 4.77, p = 0.0605; miR-199a-5p: F(1,8) = 2.28, p = 0.1697; miR-199a-5p × cisplatin interaction: F(1,8) = 0.67, p = 0.4378), despite a trend towards higher cleaved caspase-3 after cisplatin. Together with the fluorescence-based caspase-3/7 assay, these data suggest that image-based single-cell readouts captured apoptotic responses that were less evident in bulk cleaved caspase-3 measurements, particularly in the resistant context.

Taken together, these in vitro findings indicate that miR-199a-5p modulates cisplatin response in a context-dependent manner. In A2780 cells, miR-199a-5p was associated with basal XIAP repression and facilitated cisplatin-associated loss of metabolic viability and apoptotic activation. In A2780cis cells, by contrast, its effects were most evident in image-based caspase-3/7 activation and treatment-dependent XIAP regulation, without a corresponding reduction in bulk metabolic viability or robust cleaved caspase-3 accumulation.

### Spatial and subcellular reorganisation of the miR-199a-5p/XIAP axis in human HGSOC tissue

To determine whether the altered miR-199a-5p/XIAP relationship observed in vitro was reflected in tissue architecture, multiplex fluorescence in situ hybridisation combined with immunofluorescence was performed on FFPE sections from non-neoplastic ovaries and high-grade serous ovarian carcinoma (HGSOC) samples (Fig. 6). In non-neoplastic ovarian tissue, XIAP staining was intense and predominantly confined to glandular epithelial structures, whereas miR-199a-5p signal was weak and largely restricted to stromal regions (Fig. 6A). By contrast, HGSOC sections showed widespread miR-199a-5p signal across epithelial and stromal compartments, while XIAP remained detectable but displayed a more heterogeneous distribution (Fig. 6B). This redistribution was also evident at intermediate magnification: control tissue showed epithelial enrichment of XIAP with minimal overlap with miR-199a-5p (Fig. 6C, upper panel), whereas tumour tissue contained multiple cellular domains in which both signals occupied overlapping territories, compatible with partial spatial co-localisation (Fig. 6C, lower panel). Higher-magnification imaging further showed that HGSOC differed from non-neoplastic tissue not only in marker distribution but also in miR-199a-5p subcellular localisation. In control ovary, XIAP was predominantly cytoplasmic in epithelial cells, while miR-199a-5p remained faint and peripherally distributed (Fig. 6D, left). In HGSOC cells, miR-199a-5p displayed broader cytoplasmic staining together with prominent perinuclear and occasional nuclear-associated signal, a pattern absent from control tissue (Fig. 6D, right). This nuclear-associated signal was confirmed at higher magnification in multiple tumour cells (Fig. 6E), indicating tumour-associated redistribution of miR-199a-5p at both tissue and subcellular levels.

**Figure 6.**
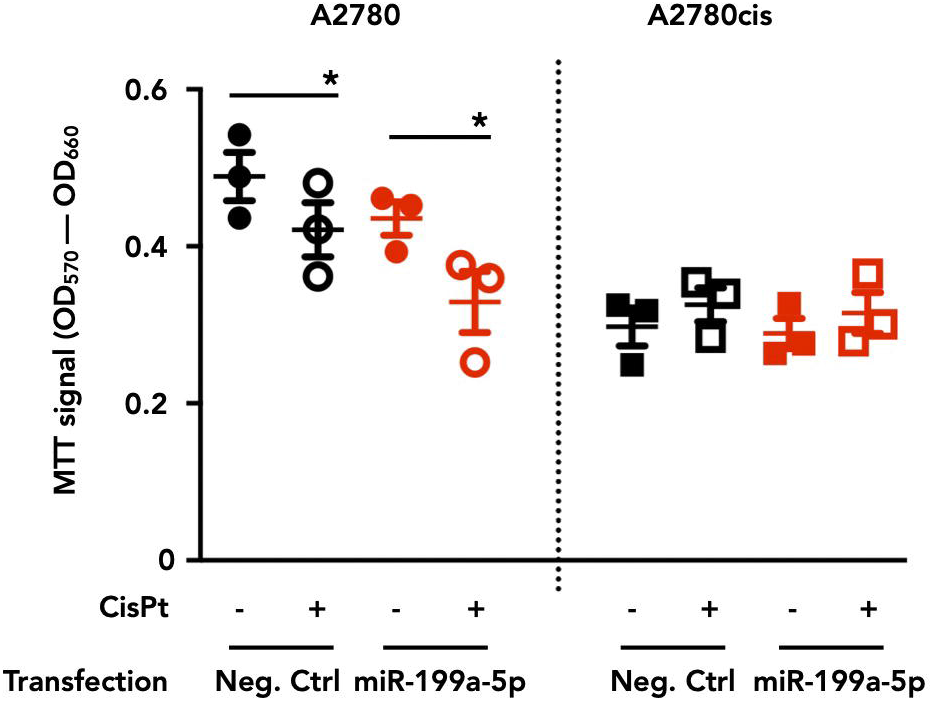
Spatial and subcellular localisation of miR-199a-5p and XIAP in non-neoplastic ovary and HGSOC. Representative multiplex fluorescence in situ hybridisation (FISH) and immunofluorescence (IF) images of formalin-fixed paraffin-embedded (FFPE) sections from non-neoplastic human ovarian tissue and high-grade serous ovarian carcinoma (HGSOC). (A, B) Low-magnification overview images of non-neoplastic ovary (A) and HGSOC (B) showing miR-199a-5p (red), XIAP (green) and nuclei (DAPI, blue). Red boxes indicate regions selected for higher-magnification analysis. (C, D) Intermediate- (C) and high-magnification (D) views of the selected regions. For each condition, panels show merged signal (left), miR-199a-5p/DAPI (middle) and XIAP/DAPI (right). (E) Digitally enlarged views of the high-magnification fields shown in (D), highlighting the subcellular distribution of miR-199a-5p in non-neoplastic ovary (left) and HGSOC (right). Arrows indicate cells showing nuclear-associated signal. Scale bars, 100 μm in (A, B), 50 μm in (C, D), and 20 μm in (E).

Quantitative colocalisation analysis revealed tumour-associated changes in the spatial relationship between miR-199a-5p and XIAP, particularly in Manders’ M1 and intensity-weighted K indices (Supplementary Table 1). Pearson’s correlation coefficient showed a modest, non-significant increase in tumours relative to controls (0.541 ± 0.142 vs. 0.464 ± 0.114; p = 0.2149). Manders’ M1 was significantly higher in tumours than in controls (0.574 ± 0.137 vs. 0.493 ± 0.062; p = 0.0220), whereas M2 remained unchanged (0.566 ± 0.191 vs. 0.510 ± 0.141; p = 0.4893). The overlap coefficient was similarly high in both groups (0.926 ± 0.025 in tumours and 0.933 ± 0.035 in controls), indicating that simple overlap alone did not capture the main tissue-associated difference.

Intensity-weighted descriptors provided stronger discrimination between groups. Control tissues showed lower K1 and higher K2 values, whereas tumour samples displayed the opposite pattern, with increased K1 (median 1.677 vs. 0.484; p = 0.000022) and reduced K2 (median 0.512 vs. 1.822; p = 0.000100). Among the metrics analysed, M1, K1 and K2 provided the clearest quantitative separation between non-neoplastic and tumour tissues, whereas Pearson’s correlation, M2 and the overlap coefficient helped delimit the effect by showing that the spatial phenotype was not driven by a uniform increase across all co-localisation parameters. These results indicate selective spatial reorganisation rather than a uniform increase in co-localisation. Together, these data identify a tumour-associated alteration of the miR-199a-5p/XIAP axis in HGSOC, characterised by loss of the spatial segregation observed in non-neoplastic ovary, increased XIAP-associated overlap with miR-199a-5p, redistribution of signal intensity within co-localised regions and altered subcellular localisation of miR-199a-5p.

## DISCUSSION

Ovarian cancer mortality remains closely linked to late-stage diagnosis and the emergence of platinum resistance [20,21]. Platinum resistance is not a single biological event, but a heterogeneous state involving DNA damage repair, survival signalling, metabolic adaptation, tumour microenvironmental cues and impaired apoptotic execution [1,5,7]. Within this framework, XIAP and other inhibitor-of-apoptosis proteins have been implicated in reduced chemotherapy-induced cell death [22], while deregulated miRNA networks have been associated with ovarian cancer progression and treatment response [8,23]. The miR-199 family is particularly relevant because it has been linked to tumour-suppressive functions and chemotherapy response in ovarian cancer [24,25]. Here, we investigated whether the miR-199a-5p/XIAP relationship remains functionally preserved across cisplatin-sensitive and resistant states, and whether this axis is spatially reorganised in human HGSOC tissue.

Our previous work identified XIAP as a direct miR-199a-5p target [13]. In the present study, STarMir and RNAfold supported an accessible and energetically favourable miR-199a-5p binding site within the XIAP 3′UTR, conserved across species. This agrees with evidence that many mammalian miRNA target sites are evolutionarily maintained [26], that RNA structure and target-site accessibility influence miRNA-mediated repression [14,27], and that target-site architecture contributes to intracellular targeting efficacy [46]. Thus, the central question was not whether XIAP can be targeted by miR-199a-5p, but whether this interaction retains the same functional meaning in different ovarian cancer contexts.

In cisplatin-sensitive A2780 cells, miR-199a-5p retained basal repression of XIAP, consistent with its validated role as a direct XIAP regulator [13]. This places miR-199a-5p within a regulatory context already relevant to platinum-induced apoptosis, since XIAP modulates Akt activity and caspase-3-dependent cleavage during cisplatin-induced cell death [30], and IAP expression has been associated with reduced cisplatin responsiveness in ovarian cancer models [31]. The reduction of XIAP after cisplatin exposure is also consistent with evidence that sensitisation to cisplatin can involve XIAP destabilisation or degradation, as described for HtrA1-mediated sensitisation [32]. However, the combined effect of miR-199a-5p and cisplatin was more compatible with additive facilitation than with strong synergy. This supports a model in which miR-199a-5p lowers the apoptotic threshold in sensitive cells while remaining aligned with XIAP repression. Such behaviour fits threshold-based models of miRNA action, where modest repression of key regulatory nodes can tune pathway responsiveness without producing binary effects across all readouts [33].

The resistant A2780cis model changed this interpretation. These cells showed reduced basal miR-199a-5p without a significant increase in XIAP mRNA or bulk XIAP protein. This agrees with ovarian cancer studies placing the miR-199 family within platinum-response networks, including HIF-1α-dependent regulation of cisplatin resistance [25], mTOR-associated restoration of cisplatin sensitivity [28] and JAG1-Notch1 activation following epigenetic silencing of miR-199b-5p [29]. However, our data refine this framework by showing that reduced miR-199a-5p is not necessarily accompanied by a reciprocal basal increase in XIAP. Therefore, the resistant state cannot be reduced to a simple “low miRNA-high target” model.

This decoupling became more apparent under treatment. In A2780cis cells, miR-199a-5p enhanced caspase-3/7 activation without producing a corresponding reduction in XIAP protein levels. This contrasts with A2780 cells, where basal XIAP repression was preserved and cisplatin-associated responses remained more closely aligned with apoptotic output. The result suggests that miR-199a-5p remains biologically active in resistant cells, but that its activity is no longer adequately represented by XIAP abundance alone. This interpretation is plausible because miR-199a has been connected to several survival-associated pathways, including HIF-1α-dependent cisplatin resistance [25], IKKβ-mediated regulation of NF-κB activity [34] and mTOR-dependent modulation of cisplatin sensitivity [28]. In resistant cells, miR-199a-5p may therefore distribute its effects across a broader stress-response network, allowing apoptotic responsiveness to change even when XIAP does not decrease proportionally at population level.

Stress-dependent changes in miRNA function provide an additional explanation for this non-linear behaviour. Cellular stress can relieve miRNA-mediated translational repression [35], and specific conditions can switch some miRNAs from repression to translational activation [36]. Although we did not directly test either mechanism, these studies show that treatment exposure can alter the output of a miRNA-target interaction. In addition, endogenous miRNA and target concentrations influence susceptibility to miRNA-mediated repression [47]. Thus, the absence of XIAP downregulation in A2780cis cells should not be interpreted as loss of miR-199a-5p activity, but as evidence that its regulatory output is redistributed in the resistant state.

The divergence between apoptosis assays supports this view. Western blotting detected only modest changes in cleaved caspase-3, whereas single-cell imaging showed clearer caspase-3/7 activation, particularly in A2780cis cells. This difference is biologically relevant because bulk assays can obscure responsive subpopulations when treatment effects are heterogeneous [37]. Recent single-cell and spatial transcriptomic analysis in HGSOC identified a platinum-resistant epithelial subcluster overexpressing TACSTD2 [50], further supporting cellular-resolution approaches for resistant disease. Our findings therefore suggest that miR-199a-5p can modulate apoptotic competence in resistant cells, but that this effect is more evident at cell level than through bulk protein accumulation alone.

The tissue data extend this model into native tumour architecture. In non-neoplastic ovary, miR-199a-5p signal was relatively stromal, whereas XIAP enrichment was mainly epithelial, suggesting spatial separation between regulatory miRNA signal and target-associated protein signal. This is consistent with evidence that stromal miRNA programmes are biologically active in ovarian cancer, including miRNA-driven reprogramming of normal fibroblasts into cancer-associated fibroblasts [38]. In HGSOC, this compartmental organisation was altered, with broader miR-199a-5p/XIAP overlap, redistribution of signal intensity and inversion of K1/K2 values accompanied by increased M1. These findings indicate a changed spatial relationship rather than a simple increase in either marker.

This spatial reorganisation is consistent with recent ovarian cancer studies showing that tissue architecture shapes tumour behaviour. Spatial transcriptomics has identified discrete tumour microenvironments and autocrine loops within ovarian cancer subclones [3,39], while artificial intelligence-guided spatial pathology has shown that prognostic regions in HGSOC may differ molecularly despite similar histology [4]. At the tumour-stroma interface, heterogeneous ligand-receptor programmes have been associated with long-term ovarian cancer survival [48], and spatial proteomic profiling has linked platinum-refractory HGSOC to coordinated AKT and WNT activity within an immunosuppressive tumour microenvironment [49]. These findings support interpreting the miR-199a-5p/XIAP axis within native tumour architecture rather than as a two-component interaction detached from spatial context.

However, increased overlap between miR-199a-5p and XIAP should not be interpreted as proof of effective repression. Colocalisation analysis measures spatial proximity, but proximity does not establish functional interaction [41]. This distinction is critical because HGSOC showed increased overlap while XIAP remained prominent within colocalised regions. Threshold-based models show that miRNAs can generate non-linear target responses [33], endogenous miRNA:target ratios determine susceptibility to miRNA-mediated competition [40,47], and target-site features influence repression efficacy [46]. Therefore, increased miR-199a-5p/XIAP overlap in HGSOC may reflect spatial coexistence under altered stoichiometric or compartmental constraints, rather than efficient repression of XIAP.

One possible explanation is that local XIAP target availability exceeds the effective repressive capacity of the miR-199a-5p pool [47]. Another is tumour-associated 3′UTR shortening, since alternative cleavage and polyadenylation can remove regulatory 3′UTR elements and activate oncogenic programmes in cancer cells [42]. We did not test XIAP 3′UTR isoform usage, AGO occupancy or direct miR-199a-5p binding to XIAP transcripts in tissue. Future work should determine whether altered spatial overlap reflects changes in target accessibility, target abundance, Argonaute loading or redistribution of miR-199a-5p away from cytoplasmic target sites.

The altered subcellular pattern of miR-199a-5p adds another layer to this interpretation. In HGSOC cells, miR-199a-5p showed prominent perinuclear and occasional nuclear-associated localisation, a pattern absent from non-neoplastic tissue. Mature miRNAs are not restricted to the cytoplasm, and nuclear import may be mediated by transport mechanisms such as importin 8-dependent miRNA trafficking [43]. Nuclear miRNAs can contribute to transcriptional and post-transcriptional regulation [9,10,44], and oncogenic nuclear miRNA activity can modulate snRNA and splicing programmes [45]. Nevertheless, our data do not prove nuclear sequestration, altered AGO loading or impaired access of miR-199a-5p to XIAP transcripts. This pattern should therefore be interpreted as a spatially resolved phenotype requiring functional validation, not as direct evidence of a nuclear mechanism.

These findings have three main implications. First, miR-199a-5p abundance alone is unlikely to predict regulatory activity, because the same miRNA remains aligned with XIAP repression in sensitive cells but becomes uncoupled from XIAP abundance in resistant cells. Second, XIAP protein abundance alone may not capture apoptotic competence when single-cell assays reveal responsive subpopulations partly obscured by bulk assays. Third, spatial organisation may influence whether miRNA and target signals coexist in a configuration compatible with repression, competition or functional uncoupling. This interpretation is consistent with the proteogenomic view that chemo-refractory HGSOC is defined by integrated molecular states rather than isolated resistance markers [51]. Future studies should test whether the miR-199a-5p/XIAP spatial phenotype correlates with platinum response in clinically annotated HGSOC cohorts and whether restoration of miR-199a-5p requires combination strategies targeting additional survival pathways. This translational direction is supported by evidence that engineered extracellular vesicles enriched with the miR-214/199a cluster can enhance chemotherapy efficacy in ovarian cancer models [11], although such approaches require validation in resistant and spatially heterogeneous disease.

Overall, our findings support a model in which the miR-199a-5p/XIAP axis is shaped by cellular state, treatment exposure and spatial organisation. In sensitive cells, miR-199a-5p remains aligned with XIAP repression and apoptotic responsiveness. In resistant cells, this alignment is reconfigured, allowing apoptotic output to remain modifiable even when XIAP abundance does not change in parallel. In human HGSOC tissue, this reconfiguration is accompanied by altered spatial overlap and subcellular redistribution. The main implication is therefore not only that miR-199a-5p targets XIAP, but that the effective activity of this interaction is dynamically organised in ovarian cancer.

## CONCLUSION

This study identifies a context-dependent reconfiguration of the miR-199a-5p/XIAP axis in ovarian cancer, in which the relationship between miR-199a-5p, XIAP and apoptotic output differs between cisplatin-sensitive and cisplatin-resistant states and is further shaped by spatial organisation in human HGSOC tissue. Our data indicate that miR-199a-5p abundance alone is insufficient to predict functional activity, highlighting the limitations of bulk expression-based approaches for assessment of miRNA-mediated regulation. By integrating functional assays with spatially resolved analysis, this work supports a framework in which miRNA activity is determined by regulatory context rather than expression level alone. These findings improve interpretation of miRNA-mediated phenotypes in ovarian cancer and support further investigation of spatially organised regulatory vulnerabilities associated with chemoresistance.

## Additional information

### Ethics approval and consent to participate

This study was approved by the Clinical Research Ethics Committee of the Complejo Hospitalario de Toledo (CEIm No. 968). Written informed consent was obtained from all participants prior to inclusion in the study. All procedures involving human participants were performed in accordance with the ethical standards of the institutional research committee and with the Declaration of Helsinki.

### Consent for publication

Not applicable.

### Availability of data and materials

The datasets generated and/or analysed during the current study are available from the corresponding author on reasonable request.

### Authors’ contributions

T.M.-G. conceived and designed the study, acquired and analysed data, interpreted the results, drafted the original manuscript, and revised the final version. D.R. and M.N.-D. contributed to data analysis and critically revised the manuscript. M.P.K., L.G.G., M.G.P. and B.S. contributed to patient recruitment, sample provision, clinical data collection, clinical interpretation, and critical revision of the manuscript. G.I. contributed to computational analysis, information systems, data processing, AI-based methodological development, and critical revision of the manuscript. R.M.M. conceived and designed the study, interpreted the results, supervised the study, and critically revised the manuscript. All authors approved the final version of the manuscript and agree to be accountable for all aspects of the work.

## Acknowledgements

The authors are deeply grateful to Dr. J.A. Rodríguez Alfaro and Dr. J. Mazarío from the Microscopy and Image Analysis Core Facilities of the National Hospital for Paraplegics, Toledo, for their outstanding technical support and expertise in microscopy and image analysis, which substantially contributed to this work.

