## Supplementary Information for "Context-dependent reconfiguration and spatial reorganisation of the miR-199a-5p/XIAP axis in high-grade serous ovarian cancer"


1. **Supplementary Methods**
    a. In silico miRNA–mRNA interaction analysis
    b. Patient samples
    c. Cell culture
    d. Transfection and cisplatin treatment
    e. RNA extraction
    f. Quantification by RT–qPCR
    g. Protein extraction and Western blot analysis
    h. Fluorescence in situ hybridisation combined with immunofluorescence in human ovarian tissue
    i. Fluorescence in situ hybridisation combined with immunofluorescence in cultured ovarian cancer cells
    j. Cell viability and cell death assays
    k. Image analysis
    l. Colocalisation analysis
    m. Statistical analysis

2. **Supplementary Figures**
    a. Supplementary Fig. S1. Cross-species computational support for conserved miR-199a-5p binding to the XIAP 3′UTR.
    b. Supplementary Fig. S2. Cisplatin sensitivity of A2780 and A2780cis ovarian cancer cells.
    c. Supplementary Fig. S3. Effects of cisplatin and miR-199a-5p on cleaved caspase-3 levels in ovarian cancer cells.

3. **Supplementary Tables**

    **Supplementary Table S1.** Quantitative colocalisation analysis of miR-199a-5p and XIAP signals in non-neoplastic ovarian tissue and high-grade serous ovarian carcinoma.

    **Supplementary Table S2.** Culture conditions for ovarian cancer cell lines used in this study.

    **Supplementary Table S3.** Primary and secondary antibodies used in this study.

    **Supplementary Table S4.** DIG-labelled LNA probe sequences used for miR-199a-5p FISH analysis.


### SUPPLEMENTARY METHODS

#### In silico miRNA–mRNA interaction analysis

The accessibility of the miR-199a-5p binding site within the human XIAP 3′ untranslated region (3′UTR) was evaluated using the STarMir web server (Wadsworth Center, New York, NY, USA; accessed 5 November 2023) [14]. The mature hsa-miR-199a-5p sequence was used as query and the human XIAP 3′UTR sequence was entered manually. Predictions were generated using the human V-CLIP-based model, with species set to *Homo sapiens* and the target region restricted to the 3′UTR. Output parameters included hybridisation energy (ΔG_hybrid), seed-pairing configuration, site accessibility scores and the composite LogitProb value. Local RNA secondary structure surrounding the predicted binding site was analysed using RNAfold from the ViennaRNA Package (Institute for Theoretical Chemistry, University of Vienna, Vienna, Austria). Human and murine XIAP 3′UTR sequences were examined under identical conditions. Minimum free-energy and centroid structures were used to assess accessibility of the predicted binding region.

**Patient samples**

Formalin-fixed paraffin-embedded ovarian tissue samples were obtained from the Department of Obstetrics and Gynecology, Hospital Universitario de Torrejón (Madrid, Spain). The cohort included 10 high-grade serous ovarian carcinoma samples and 10 non-neoplastic ovarian controls derived from oophorectomy or hysterectomy performed for benign gynaecological conditions. Representative haematoxylin and eosin-stained sections were reviewed by a certified pathologist to confirm histopathological diagnosis. For each FFPE block, a 3 mm punch biopsy was taken from the selected region of interest. Samples were anonymised, numerically coded and analysed in a blinded manner.

**Cell culture**

Human ovarian carcinoma A2780 cells (ECACC 93112519; RRID: CVCL_0134) and their cisplatin-resistant counterpart A2780cis (ECACC 93112517; RRID: CVCL_0135) were obtained from the Cell Bank of the Centro de Instrumentación Científica, University of Granada (Granada, Spain) (Supplementary Table S1). Cell lines were authenticated by short tandem repeat profiling and confirmed to be free of bacterial, fungal and mycoplasma contamination. Cells were maintained under biosafety level 2 conditions.

A2780 cells were cultured in RPMI-1640 supplemented with 10% fetal bovine serum and 2 mM L-glutamine. A2780cis cells were maintained in the same medium supplemented with 1 μM cisplatin to preserve the resistant phenotype (Supplementary Table S1). Cells were incubated at 37 °C in a humidified atmosphere containing 5% CO2. For passaging, cells at 70–80% confluence were detached using 0.25% trypsin–EDTA and reseeded at ratios of 1:3–1:6 for A2780 and 1:5–1:20 for A2780cis. After thawing, A2780cis cells were maintained in cisplatin-free medium for 24–48 h before re-exposure to cisplatin.

**Transfection and cisplatin treatment**

Transient transfections were performed using DharmaFECT-4 reagent (Dharmacon, Horizon Discovery, Waterbeach, UK) according to the manufacturer's instructions. Cells were transfected at approximately 75–80% confluence for 24 h with 50 nM miRIDIAN hsa-miR-199a-5p mimic (miRBase accession: MIMAT0000231; sequence: 5′-CCCAGUGUUCAGACUCCUGUUC-3′) or a non-targeting negative control mimic (cel-miR-67-3p; miRBase accession: MIMAT0000039; sequence: 5′-UCACAACCUCCUAGAAAGAGUAGA-3′). Following transfection, cells were treated with 12.5 µM cisplatin for 24 h under standard culture conditions. Control cells were processed in parallel and maintained in cisplatin-free medium.

**RNA extraction**

Total RNA was isolated from confluent untreated A2780 and A2780cis cultures using QIAzol Lysis Reagent (Qiagen, Hilden, Germany), followed by purification with the miRNeasy Mini Kit (Qiagen), according to the manufacturer's instructions. RNA concentration and purity were assessed using a NanoDrop ND-1000 spectrophotometer (Thermo Fisher Scientific).

**Quantification by RT–qPCR**

For miR-199a-5p quantification, 10 ng of total RNA was reverse transcribed and amplified using a TaqMan MicroRNA assay (Applied Biosystems; assay ID 000498), with U6 small nuclear RNA (assay ID 001973) as endogenous control. For XIAP mRNA quantification, 1 µg of total RNA was treated with DNase I (Roche) and reverse transcribed using Moloney murine leukaemia virus reverse transcriptase and random primers. Quantitative PCR was performed using TaqMan Universal PCR Master Mix with gene-specific probes for XIAP (Hs00180269_m1) and miR-199a-5p (000498). 18S rRNA (Hs99999901_s1) and U6 RNA (001973) were used as endogenous controls for mRNA and miRNA normalisation, respectively. Amplification was performed on a TaqMan 7900HT Fast Real-Time PCR System under standard cycling conditions (10 min at 95 °C, followed by 40 cycles of 15 s at 95 °C and 1 min at 60 °C). All experiments included three independent biological replicates and three technical replicates per condition. Relative expression was calculated using the 2^-ΔΔCt method [16], with error propagation as described by Headrick [17]. Results were expressed as fold change in A2780cis relative to A2780, including 95% confidence intervals.

**Protein extraction and Western blot analysis**

For protein analyses, A2780 and A2780cis cells were seeded in 6-well plates at 2.5 × 10^5 cells per well and cultured to approximately 80% confluence. Depending on the experimental condition, cells were left untreated or transfected with miR-199a-5p or negative control mimics for 24 h, followed, where indicated, by treatment with 12.5 µM cisplatin for an additional 24 h. Cells were harvested by mechanical detachment and lysed in RIPA buffer supplemented with cOmplete EDTA-free protease inhibitor cocktail. Lysates were incubated for 30 min at 4 °C and clarified by centrifugation at 12,000 × g for 10 min at 4 °C. Protein concentration was measured by BCA assay. Equal amounts of protein (50 µg per lane) were mixed with Laemmli sample buffer containing β-mercaptoethanol, boiled for 5 min at 100 °C, separated by SDS–PAGE and transferred onto 0.2 µm PVDF membranes. Membranes were blocked for 1 h at room temperature in 5% non-fat dry milk in TBS-T and incubated overnight at 4 °C with primary antibodies diluted in blocking solution. After washing, membranes were incubated for 90 min at room temperature with HRP-conjugated secondary antibodies. Signals were detected using chemiluminescent substrate and visualised with an ImageScanner III system. Image acquisition was performed under non-saturating conditions. Band intensities were quantified using ImageJ v1.54p. Antibody details are provided in Supplementary Table S2.

**Fluorescence in situ hybridisation combined with immunofluorescence in human ovarian tissue**

Fluorescence in situ hybridisation (FISH) for miR-199a-5p in FFPE human ovarian tissue sections was performed using an adapted RNase-free protocol based on Søe et al. [15]. All solutions were prepared using DEPC-treated water. Slides were incubated at 65 °C for 2 h, deparaffinised in xylene and rehydrated through graded ethanol before immersion in distilled water. Antigen retrieval was performed in 250 mM EDTA at 97 °C for 20 min using a PT Link system (Agilent Technologies, Santa Clara, CA, USA). Tissue digestion was carried out with pepsin solution for 30 min at 37 °C. To reduce non-specific ionic binding, sections were incubated in acetylation buffer containing triethanolamine, HCl and acetic anhydride for 10 min at room temperature. DIG-labelled LNA probes specific for miR-199a-5p or the negative control cel-miR-67 were applied to each section. Probes were denatured at 85 °C for 15 min and hybridised overnight at 37 °C. After hybridisation, slides were washed in 0.4× SSC containing 0.3% NP-40 at 75 °C for 2 min, followed by 0.2× SSC containing 0.1% NP-40 for 1 min at room temperature. Sections were blocked for 15 min at 37 °C in 5% horse serum and 1% BSA in PBS-T-DEPC and then incubated with alkaline phosphatase-conjugated anti-digoxigenin Fab fragments to detect DIG-labelled LNA probes. Following FISH detection, the same sections were processed for XIAP immunofluorescence. Sections were blocked and permeabilised in blocking solution containing normal goat serum and Triton X-100 in PBS and then incubated overnight at 4 °C with a primary anti-XIAP antibody diluted in blocking solution. After washing in PBS, sections were incubated for 2 h at room temperature with the appropriate Alexa Fluor-conjugated secondary antibody. Sections were mounted using fluorescence-compatible mounting medium containing DAPI for nuclear counterstaining. Fluorescent signals were acquired using a fully motorised Olympus IX83 microscope equipped with CellSens Dimensions software v4.1.1. Probe sequences are listed in Supplementary Table S3, and antibody details are provided in Supplementary Table S2.

**Fluorescence in situ hybridisation combined with immunofluorescence in cultured ovarian cancer cells**

FISH analysis of miR-199a-5p was performed in A2780 and A2780cis cells grown on sterile glass coverslips using an adapted RNase-free protocol with DIG-labelled LNA probes. All solutions were prepared using DEPC-treated water. Cells were washed twice with PBS-DEPC and permeabilised with PBS containing 0.1% Tween-20 for 15 min at room temperature. To reduce non-specific ionic binding, cells were incubated in freshly prepared acetylation buffer containing triethanolamine, HCl and acetic anhydride, followed by washing in PBS-DEPC. Cells were preincubated for 30 min at 65 °C in a 1:1 mixture of miRCURY LNA hybridisation buffer and DEPC-treated water. DIG-labelled LNA probes specific for miR-199a-5p or the negative control cel-miR-67 were diluted in hybridisation buffer to a final concentration of approximately 200 nM, denatured at 80 °C for 5 min and incubated with the cells for 1 h at 65 °C. Post-hybridisation washes were performed sequentially in 5×, 1× and 0.1× SSC at 65 °C, followed by a final wash in 0.1× SSC at room temperature. Samples were incubated in blocking buffer containing horse serum and BSA in PBS-T, followed by incubation with alkaline phosphatase-conjugated anti-digoxigenin Fab fragments to detect DIG-labelled LNA probes. Following FISH detection, the same coverslips were processed for XIAP immunofluorescence. Cells were blocked and permeabilised in blocking solution containing normal goat serum and Triton X-100 in PBS and incubated with a primary anti-XIAP antibody diluted in blocking solution. After washing in PBS, cells were incubated with the appropriate Alexa Fluor-conjugated secondary antibody. Coverslips were mounted on glass slides using fluorescence-compatible mounting medium containing DAPI for nuclear counterstaining. Fluorescence images were acquired using a Leica DM5000B epifluorescence microscope with a 20× objective, capturing five fields per sample. No signal was detected in negative control probe conditions or in controls processed without primary antibody. Probe sequences are listed in Supplementary Table S3, and antibody details are provided in Supplementary Table S2.

**Cell viability and cell death assays**

**MTT cell viability assay:** Cell viability was assessed using the MTT assay as a measure of mitochondrial metabolic activity. A2780 and A2780cis cells were seeded in clear 96-well plates at $2 \times 10^4$ cells per well and allowed to adhere overnight. Transfection and treatment were performed as described above, followed by exposure to 12.5 µM cisplatin or vehicle for 24 h. MTT reagent was added directly to the culture medium at a final concentration of 0.5 mg/mL and cells were incubated for 3 h at 37 °C protected from light. Formazan crystals were solubilised with HCl : isopropanol solution (1:500). Absorbance was measured at 570 nm with background correction at 660 nm using an Infinite M200 Pro microplate reader (Tecan Group Ltd.,

Männedorf, Switzerland). Cell viability was expressed as normalised absorbance relative to the untreated negative control condition.

**Propidium iodide staining assay:** Late cell death was assessed by propidium iodide staining as a measure of plasma membrane integrity. Following transfection and treatment, A2780 and A2780cis cells grown on sterile glass coverslips were incubated for 30 min at 37 °C in warm phosphate-buffered saline containing 0.4 μg/mL propidium iodide, protected from light. Cells were washed twice with PBS and mounted using DAKO Fluorescence Mounting Medium. Fluorescence images were acquired using a Leica DM5000B epifluorescence microscope equipped with a 20× objective and a Leica DFC 3000 G digital camera. The percentage of PI-positive cells relative to the total number of nuclei was used as an index of late irreversible cell death.

**Caspase-3/7 activity assay:** Effector caspase activity was assessed at single-cell level using the CellEvent Caspase-3/7 Green Detection Reagent. Following transfection and treatment, A2780 and A2780cis cells grown on sterile glass coverslips were incubated for 30 min at 37 °C in warm phosphate-buffered saline containing 2.5 μM reagent and 10% fetal bovine serum, protected from light. After incubation, the reagent was removed and cells were fixed with 4% paraformaldehyde for 15 min at room temperature. Cells were washed three times in PBS, with DAPI added during the first wash for nuclear counterstaining. Samples were mounted using DAKO Fluorescence Mounting Medium. Fluorescence images were acquired using a Leica DM5000B epifluorescence microscope equipped with a 20× objective and a Leica DFC 3000 G digital camera. The proportion of caspase-3/7-positive cells relative to the total number of nuclei was used as an index of effector caspase activation.

### Image analysis

#### In vitro experiments

Quantitative image analysis of propidium iodide staining, caspase-3/7 activity, and basal XIAP and miR-199a-5p fluorescence was performed using QuPath v0.4.3. Nuclei were detected from DAPI fluorescence using the Cell detection algorithm with the following settings: detection channel 3, background radius 15 px, median filter radius 2 px, sigma 8 px, minimum area 10 px^2, maximum area 5000 px^2, threshold 1000, and cell expansion 5 μm. The Make measurements option was enabled. XIAP immunofluorescence and caspase-3/7 activity were quantified in the FITC/Alexa 488 channel, whereas propidium iodide staining was analysed in the red channel. A total of 556 A2780 cells and 564 A2780cis cells were analysed for basal fluorescence measurements across three independent experiments. For PI and caspase-3/7 assays, approximately 1500 cells per condition were analysed across five non-overlapping fields per replicate in four independent experiments. Mean fluorescence intensity values were exported for downstream statistical analysis.

##### Human tissue FISH/IF analysis

FISH/IF images were acquired using a fully motorised Olympus IX83 microscope with CellSens Dimensions software v4.1.1. Image analysis was performed in QuPath v0.4.3 using a custom macro. Ten regions of interest (750 × 750 µm) were defined per sample to ensure representative tissue coverage. Nuclei were automatically detected based on DAPI staining, and fluorescence intensities were quantified per detected object in the FITC (XIAP) and Cy5 (miR-199a-5p) channels. ROI-level CSV files were exported for downstream analysis.

#### Colocalisation analysis

Colocalisation metrics were computed in a blinded manner using R (≥4.2) within the tidyverse framework. Channel-specific thresholds were defined using the 25th percentile of per-cell intensity distributions. For each ROI, Pearson correlation coefficients and Manders' overlap coefficients (M1 and M2) were calculated from per-cell mean intensities. Cross-correlation descriptors exported from QuPath (CCF_MAX, CCF_MIN, K1, K2 and FWHM) were also incorporated. ROI-level data were aggregated to generate a single mean value per image by averaging across the 10 ROIs. Both ROI-level and image-level datasets were exported for statistical analysis, with group labels masked during processing to preserve blinding.

#### Statistical analysis

Statistical analysis was performed using GraphPad Prism and R (≥4.2). Linear mixed-effects models were used as the primary analytical framework for functional assays, including MTT, PI, caspase-3/7 activation and XIAP protein levels under treatment. In these models, transfection status, cisplatin treatment and their interaction were included as fixed effects, while experimental day was included as a random intercept. Fixed effects were evaluated using Satterthwaite's approximation, as implemented in the *lmerTest* package. Post hoc pairwise comparisons were adjusted using Tukey's method where appropriate.

For two-group comparisons under basal conditions, Student's *t*-tests were applied to RT–qPCR measurements, whereas basal FISH/IF fluorescence intensities were analysed using Mann–Whitney *U* tests. A paired one-tailed *t*-test was used for densitometric analysis of basal XIAP immunoblots following miR-199a-5p transfection, based on a pre-specified directional hypothesis of XIAP reduction. For human tissue analyses and single-cell intensity distributions that did not meet normality or homoscedasticity assumptions, non-parametric or variance-robust tests were used. Specifically, Mann–Whitney *U* tests were used for M1, K1 and FWHM metrics, whereas Welch's *t*-tests were used for Pearson correlation, M2, CCF_MAX, CCF_MIN and K2 parameters. Principal component analysis was applied to explore multivariate structure within the colocalisation dataset.

Data are presented as mean ± standard error of the mean (SEM) calculated from independent experiments unless otherwise stated. For image-based assays, measurements from multiple fields and cells within each experiment were aggregated at the experimental level, and each independent experiment was treated as one biological replicate for statistical inference. All tests were two-sided unless otherwise specified. Statistical significance was achieved when $P < 0.05$.

### SUPPLEMENTARY FIGURES

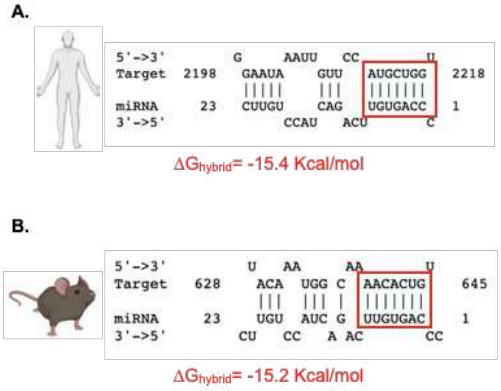


C.

| miRNA | Site Position | LogitProb† | Score | $\Delta G_{hybrid}$ |
|---|---|---|---|---|
| hsa-miR-199a-5p | 2198-2218 | 0.543 | 0.234 | -15.4 |

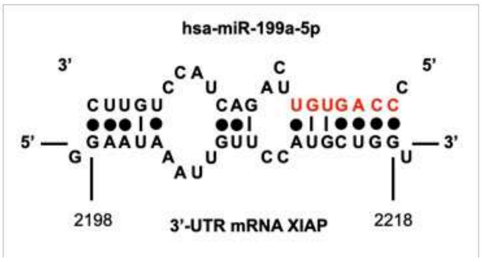


**Supplementary Fig. S1. Cross-species computational support for conserved miR-199a-5p binding to the XIAP 3′UTR. A.** Predicted duplex formed between hsa-miR-199a-5p and the human XIAP 3′ untranslated region (3′UTR; positions 2198–2218), showing canonical seed pairing and extended base complementarity. **B.** Predicted conserved interaction between miR-199a-5p and the murine XIAP orthologue (positions 628–645), also showing canonical seed pairing and comparable hybridisation energy ($\Delta G\_hybrid = -15.2$ kcal/mol). **C.** Summary of STarMir predictions for the human XIAP target site. Site_Position indicates the nucleotide coordinates of the predicted binding region within the human XIAP 3′UTR. LogitProb represents the probability of a functional miRNA-binding site, as estimated by a logistic regression model trained on CLIP-derived datasets; values >0.5 were considered supportive of binding. Score integrates multiple determinants of interaction, including seed pairing, 3′ pairing, target accessibility, AU-rich flanking context and hybridisation energy. More negative ΔG_hybrid values indicate greater thermodynamic stability of duplex formation. **D.** RNA secondary structure model of the human XIAP 3′UTR surrounding the predicted miR-199a-5p binding site, generated using RNAfold. The predicted miRNA-binding region is highlighted, and dots indicate base-pairing interactions within the local RNA structure.

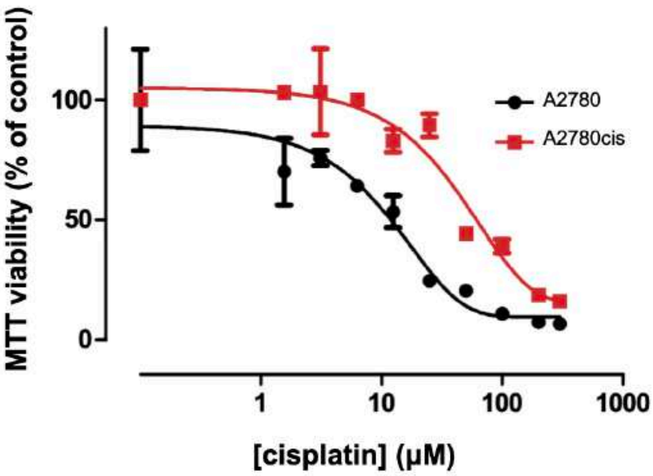


**Supplementary Fig. S2. Cisplatin sensitivity of A2780 and A2780cis ovarian cancer cells.** Dose–response curves were generated from MTT assays following 72 h exposure to increasing concentrations of cisplatin. $EC_{50}$ values were calculated to compare cisplatin sensitivity between the two cell lines. A2780cis cells showed a higher cisplatin $EC_{50}$ than parental A2780 cells (44.56 μM vs. 12.16 μM), confirming the resistant phenotype. Data represent mean ± SEM from three independent experiments performed in triplicate.

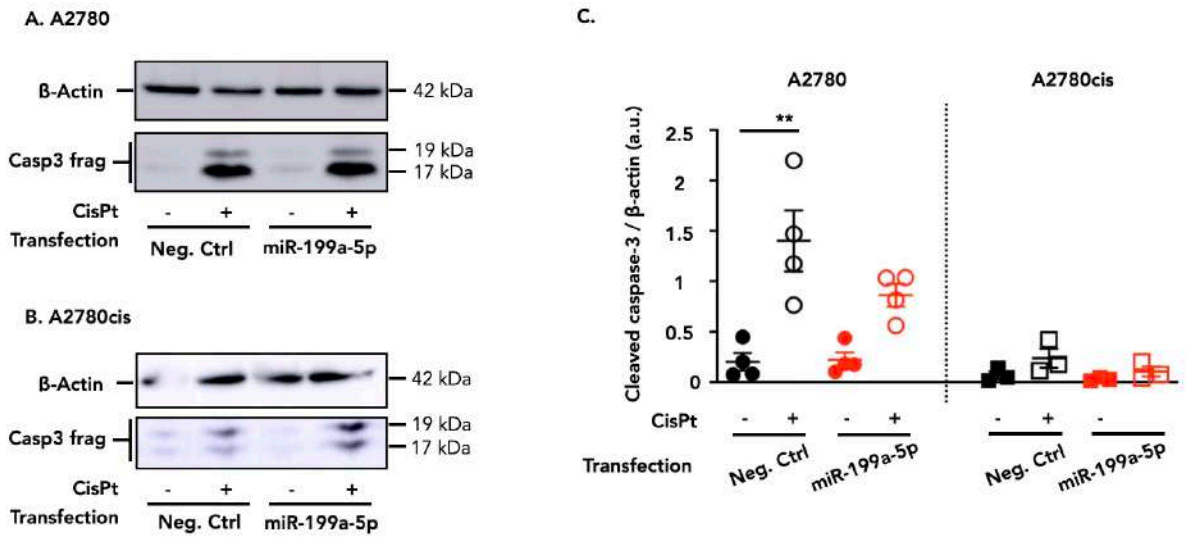


**Supplementary Fig. S3. Effects of cisplatin and miR-199a-5p on cleaved caspase-3 levels in ovarian cancer cells. A and B.** Representative western blot analysis of cleaved caspase-3 (Casp3 frag) in whole-cell lysates from A2780 (A) and A2780cis (B) ovarian cancer cells transfected with either a negative control mimic or a miR-199a-5p mimic, and treated with or without cisplatin (CisPt; 12.5 µM, 24 h). β-Actin was used as a loading control. **C.** Densitometric quantification of cleaved caspase-3 normalised to β-actin. A2780 cells are shown on the left (circles; n = 4) and A2780cis cells on the right (squares; n = 3). Each point represents one independent experiment, and bars indicate mean ± SEM. Statistical analysis was performed using a linear mixed-effects model with transfection, cisplatin treatment and their interaction as fixed effects, and experimental day as a random intercept, followed by Tukey-adjusted post hoc comparisons. **$p < 0.01$.

### SUPPLEMENTARY TABLES

**Supplementary Table S1.** Quantitative colocalisation analysis of miR-199a-5p and XIAP signals in non-neoplastic ovarian tissue and high-grade serous ovarian carcinoma.

| Variable | Control | Tumor | Test | p_value |
|---|---|---|---|---|
| **Pearson** | 0.464 ± 0.114 | 0.541 ± 0.142 | Welch t-test | 0.214901 |
| **M1** | **0.493 ± 0.062** | **0.574 ± 0.137** | **Mann–Whitney U** | **0.022018** |
| **M2** | 0.510 ± 0.141 | 0.566 ± 0.191 | Welch t-test | 0.489253 |
| **Overlap coefficient** | 0.933 ± 0.035 | 0.926 ± 0.025 | | |
| **CCF_MAX** | 0.456 ± 0.115 | 0.534 ± 0.141 | Welch t-test | 0.211654 |
| **CCF_MIN** | 0.296 ± 0.101 | 0.371 ± 0.130 | Welch t-test | 0.181979 |
| **K1** | **0.484 (median)** | **1.677 (median)** | **Mann–Whitney U** | **0.000022** |
| **K2** | **1.822 (median)** | **0.512 (median)** | **Mann–Whitney U** | **0.000100** |
| **FWHM** | 20.506 (median) | 20.235 (median) | Mann–Whitney U | 0.660720 |

**Supplementary Table S2:** Culture media and supplements used for each cell lines

| Cell line | Media | Supplements |
|---|---|---|
| A2780[1] - (ECACC 93112519; RRID: CVCL_0134) | RPMI-1640[2] | 10% fetal bovine serum[2]; 2 mM L-glutamine[2] |
| A2780cis[1] - (ECACC 93112517; RRID: CVCL_0135) | RPMI-1640[2] | 10% fetal bovine serum[2]; 2 mM L-glutamine[2]; 1 μM cisplatin[3] |

[1] Cell Bank of the Centro de Instrumentación Científica (CIC), University of Granada (Granada, Spain).
[2] Gibco, Thermo Fisher Scientific, Waltham, MA, USA.
[3] Merck KGaA, Darmstadt, Germany.

**Supplementary Table S3.** Primary and secondary antibodies used in this study.

| Antibody | Reference |
|---|---|
| **Primary antibodies for Immunoblot** | |
| anti-XIAP | BD Biosciences (Franklin Lakes, NJ, USA) Cat# 610716, RRID:AB_398039) |
| anti-β-Actin | BD Biosciences (Franklin Lakes, NJ, USA); Cat# 612656, RRID:AB_2289199 |
| **Primary antibodies for Immunofluorescence** | |
| anti-XIAP | Abcam (Cambridge, UK); Cat# ab21278; RRID: AB_446157 |
| anti-Caspase 3 fragment | Cell Signaling Technology (Danvers, MA, USA); Cat# 94530; RRID: AB_3076239 |
| **Primary antibody for FISH** | |
| Anti-digoxigenin Fab fragments, alkaline phosphatase-conjugated | Sigma-Aldrich Cat# 11093274910, RRID:AB_2734716 |
| **Secondary antibodies for Immunoblot** | |
| HRP-conjugated goat anti-rabbit | Cell Signaling Technology (Danvers, MA, USA); Cat# 7074, RRID:AB_2099233 |
| HRP-conjugated goat anti-mouse | Cell Signaling Technology; Cat# 7076, RRID:AB_330924 |
| **Secondary antibodies for Immunofluorescence** | |
| Alexa Fluor 488 goat anti-mouse, highly cross-adsorbed | Molecular Probes (Eugene, OR, USA); Cat# A11029, RRID:AB_138404 |

**Supplementary Table S4.** DIG-labelled LNA probe sequences used for miR-199a-5p FISH analysis. Probes were designed according to Søe et al. [15] and contain 2′-O-methyl RNA nucleotides, indicated by square brackets, and locked nucleic acid (LNA) nucleotides, indicated by braces.

| Probe | Sequence |
|---|---|
| **Negative Control[1]** | 5′-DIG-{G}[T]{GU}[A]{AC}[A]{CG}[T]{CU}[A]{UA}[C]{GC}[C]{CA}-3′ |
| **miR-199a-5p[1]** | 5′-DIG-{G}[AA]{C}[AG]{G}[UA]{G}[UC]{T}[GA]{A}[CA]{C}[UG]{GG}-3′ |

[1]Eurogentec, Seraing, Belgium.
